# High-speed compressed-sensing fluorescence lifetime imaging microscopy of live cells

**DOI:** 10.1101/2020.07.16.205161

**Authors:** Yayao Ma, Youngjae Lee, Catherine Best-Popescu, Liang Gao

## Abstract

We present high-resolution, high-speed fluorescence lifetime imaging microscopy (FLIM) of live cells based on a compressed sensing scheme. By leveraging the compressibility of biological scenes in a specific domain, we simultaneously record the time-lapse fluorescence decay upon pulsed laser excitation within a large field of view. The resultant system, referred to as compressed FLIM, can acquire a widefield fluorescence lifetime image within a single camera exposure, eliminating the motion artifact and minimizing the photobleaching and phototoxicity. The imaging speed, limited only by the readout speed of the camera, is up to 100 Hz. We demonstrated the utility of compressed FLIM in imaging various transient dynamics at the microscopic scale.

---

Ultrafast biological dynamics occur at atomic or molecular scales. Resolving these transient phenomena requires a typical frame rate at GHz, which is beyond the bandwidth of most electronic sensors, posing a significant technical challenge on the detection system^1,2^. Ultrafast optical imaging techniques^3–7^ can provide a high temporal resolution down to a femtosecond and have established themselves indispensable tools in blur-free observation of fast biological events^6–11^.

Fluorescence lifetime imaging microscopy (FLIM)^12,13^—a powerful ultrafast imaging tool for fingerprinting molecules—has been extensively employed in a wide spectrum of biomedical applications, ranging from single molecule analysis to medical diagnosis^14,15^. Rather than imaging the time-integrated fluorescent signals, FLIM measures the time-lapse fluorescent decay after pulsed or time-modulated laser excitation. Because the lifetime of a fluorophore is dependent on its molecular environment but not on its concentration, FLIM enables a more quantitative study of molecular effects inside living organisms compared with conventional intensity-based approaches^15–20^.

To detect the fast fluorescence decay in FLIM, there are generally two strategies. Time-domain FLIM illuminates the sample with pulsed laser, followed by using time-correlated single photon counting (TCSPC) or time-gated sensors to directly measure the fluorescence decay^13,21,22^. By contrast, frequency-domain FLIM excites the sample with high-frequency modulated light and infers the fluorescence lifetime through measuring the relative phase shift in the modulated fluorescence^23,24^. Despite being quantitative, the common drawback of these approaches is their dependence on scanning and/or repetitive measurements. For example, to acquire a two-dimensional (2D) image, a confocal FLIM imager must raster scan the entire field of view (FOV), resulting in a trade-off between the FOV and frame rate. To avoid motion artifacts, the sample must remain static during data acquisition. Alternatively, widefield FLIM systems acquire spatial data in parallel. Nonetheless, to achieve high temporal resolution, they still need to temporally scan either a gated window^25,26^ or detection phase^27^, or they must use TCSPC which requires a large number of repetitive measurements to construct a temporal histogram ^28,29^. Limited by the scanning requirement, current FLIM systems operate at only a few frames per second when acquiring high-resolution images^21,30,31^. The slow frame rate thus prevents these imagers from capturing transient biological events, such as neural spiking^32^ and calcium oscillation^33^. Therefore, there is an unmet need to develop new and efficient imaging strategies for high-speed, high-resolution FLIM.

To overcome the above limitations, herein we introduced the paradigm of compressed sensing into FLIM and developed a snapshot widefield FLIM system, termed compressed FLIM, which can image fluorescence lifetime at an unprecedented speed. Our method is made possible by a unique integration of (1) compressed ultrafast photography (CUP)^5^ for data acquisition, (2) a dual-camera detection scheme for high-resolution image reconstruction^34^, and (3) computer cluster hardware and graphic processing unit (GPU) parallel computing technologies for real-time data processing. The synergistic effort enables high-resolution (500 × 400) widefield lifetime imaging at 100 frames per second (fps). To demonstrate compressed FLIM, we performed experiments sequentially on fluorescent beads and live neurons, measuring the dynamics of bead diffusion and firing of action potential, respectively.

## Results

### Compressed FLIM

Compressed FLIM operates in two steps: data acquisition and image reconstruction (both further described in Methods). Briefly, the sample is first imaged by a high-resolution fluorescence microscope. The output image is then passed to the CUP camera for time-resolved measurement. Finally, we use a GPU-accelerated compressed sensing reconstruction algorithm—two-step iterative shrinkage/thresholding (TwIST)^35^—to process the image in real time.

A compressed FLIM system (**Fig. 1**) consists of an epi-fluorescence microscope and a CUP camera. An animated video (**Movie 1**) shows the system in action. Upon excitation by a single laser pulse, the fluorescence is collected by an objective lens with a high numerical aperture (NA) and forms an intermediate image at the microscope’s side image port. A beam splitter then divides the fluorescence into two beams. The reflected light is directly captured by a reference complementary metal–oxide–semiconductor (CMOS) camera, generating a time-integrated image:

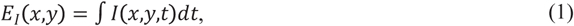

**Fig.1.**
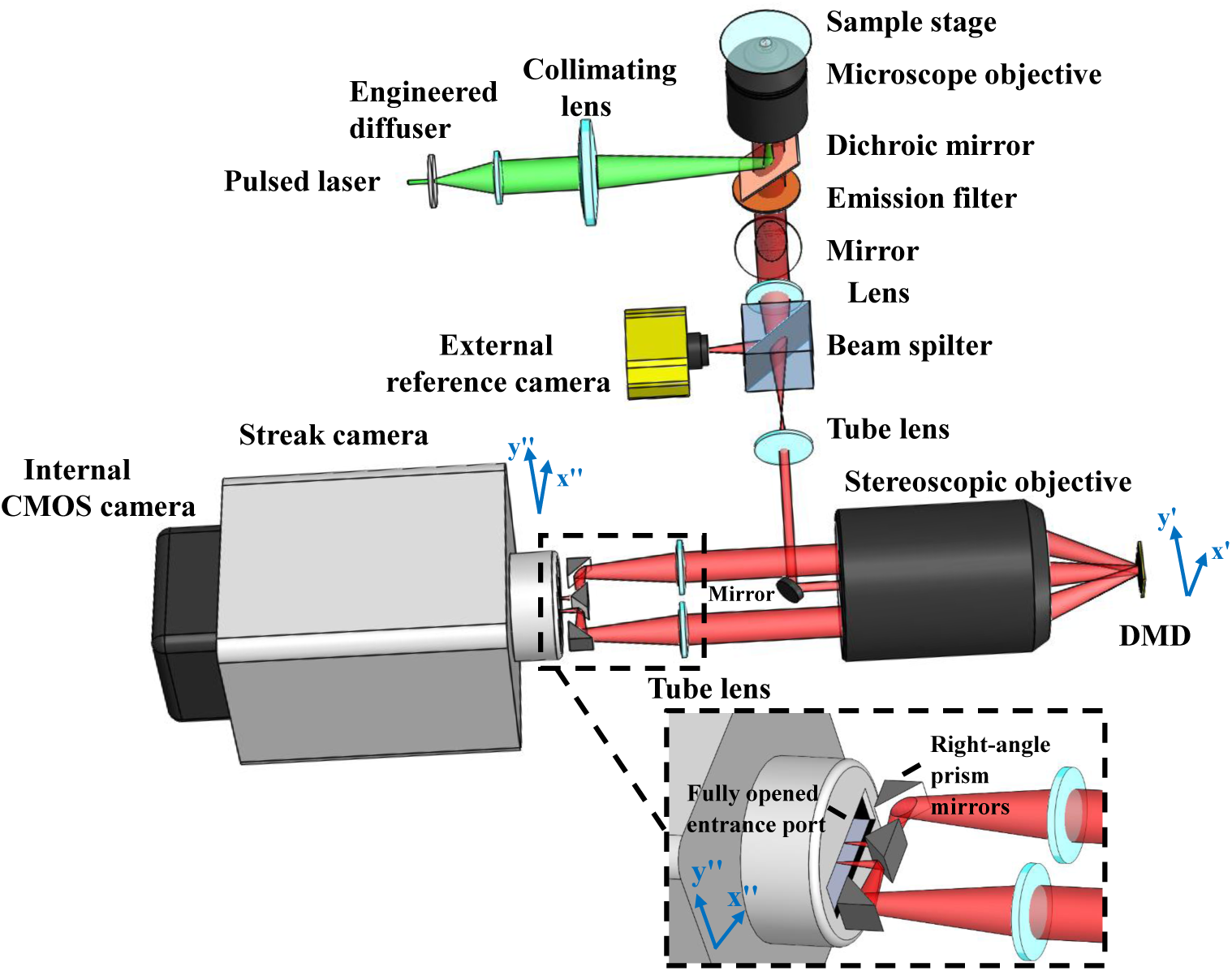
Schematic of compressed FLIM. Lower right inset: Close-up of the configuration at the streak camera’s entrance port. Light beams in two complementary encoding channels are folded by two right angle prisms before entering the fully opened entrance port of the streak camera. DMD: digital micromirror device. CMOS: complementary metal–oxide–semiconductor.

where *I*(*x,y,t*) denotes the time-lapse fluorescence decay.

The transmitted light is relayed to a digital micromirror device (DMD) through a 4*f* imaging system consisting of a tube lens, a mirror, and a stereoscope objective. A static, pseudorandom, binary pattern is displayed on the DMD to encode the image. Each encoding pixel is turned either on (tilted −12° with respect to the DMD surface norm) or off (tilted +12° with respect to the DMD surface norm) and reflects the incident fluorescence in one of the two directions. Two reflected fluorescence beams, masked with the complementary patterns, are both collected by the same stereoscope objective and enter corresponding sub-pupils at the objective’s back focal plane. The fluorescence from these two sub-pupils are then imaged by two tube lenses, folded by right-angle prism mirrors (the lower right inset in **Fig. 1**), and form two complementary channel images at the entrance port of a streak camera. To accept the encoded 2D image, this entrance port is fully opened (∼5 mm width) to its maximum, exposing the entire photocathode to the incident light. Inside the streak camera, the encoded fluorescence signals are temporally deflected along the vertical axis according to the time of arrival of incident photons. The final image is acquired by a CMOS camera—the photons are temporally integrated within the camera’s exposure time and spatially integrated within the camera’s pixel. The formation of complementary channel images can be written as:

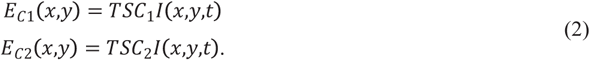

Here *S* is a temporal shearing operator, and *T* is a spatio-temporal integration operator. They describe the functions of the streak camera and CMOS camera, respectively. *C*_1_ and *C*_2_ are spatial encoding operators, depicting the complementary masks applied to two channel images, and *C*_1_ + *C*_2_ = *I*, where *I* is a matrix of ones. This complementary-encoding setup features a 100% light throughput, saving every photon in low-light conditions. Also, because there is no information lost, our encoding scheme enriches the observation and favors the compressed-sensing-based image reconstruction.

During data acquisition, we synchronize the streak camera with the reference camera in the transmission optical path. Therefore, each excitation event yields three images: one time-integrated fluorescence image, *E*_*I*_*(x,y)*, and two spatially-encoded, temporally-sheared channel images, *E*_*C*1_*(x,y)* and *E*_*C*2_*(x,y)*.

The image reconstruction of compressed FLIM is the solution of the inverse problem of the above image formation process (Eq. 1-2). Because the two complementary channel images *E*_*C*1_, *E*_*C*2_ are essentially associated with the same scene, the original fluorescence decay event can be reasonably estimated by applying a compressed sensing algorithm TwIST^35^ to the concatenated data (*E*_*C*1_, *E*_*C*2_). Additionally, to further improve the resolution, we apply the time-integrated image recorded by the reference camera, *E*_*I*_, as the spatial constraint. Finally, we fit a non-linear least-squares exponential curve to the reconstructed fluorescence decay at each spatial sampling location and produce the high-resolution fluorescence lifetime map. To reconstruct the image in real time, we implement our algorithm on GPU and computer cluster hardware.

The synergistic integration of hardware and algorithm innovations enables acquisition of high-quality microscopic fluorescence lifetime images. Operating in a snapshot format, the frame rate of compressed FLIM is limited by only the readout speed of the streak camera and up to 100 fps. The spatial resolution, determined by the numerical aperture of the objective lens, is in a submicron range, providing a resolving power to uncover the transient events inside a cell.

### Imaging fluorescent beads in flow

We demonstrated compressed FLIM in imaging fluorescent beads in flow. We mixed two types of fluorescent beads (diameters, 6 μm and 2 μm) in phosphate buffer solutions (PBS) and flowed them in a microtubing at a constant speed using a syringe pump. The ground-truth fluorescence lifetimes of these two types of fluorescent beads are 5.0 ns and 3.6 ns, respectively. We excited the beads at 532 nm and continuously imaged the fluorescence using compressed FLIM at 75 fps. As an example, the reconstructed time-lapse fluorescence decays at two beads’ locations are shown in **Fig. 2a**. We pseudo-colored the bead image based on the fitted lifetimes (**Fig. 2c**). The result indicates that compressed FLIM can differentiate these two types of beads with very close lifetimes. For comparison, the corresponding time-integrated image captured by the reference camera is shown in **Fig. 2d**.

**Fig.2.**
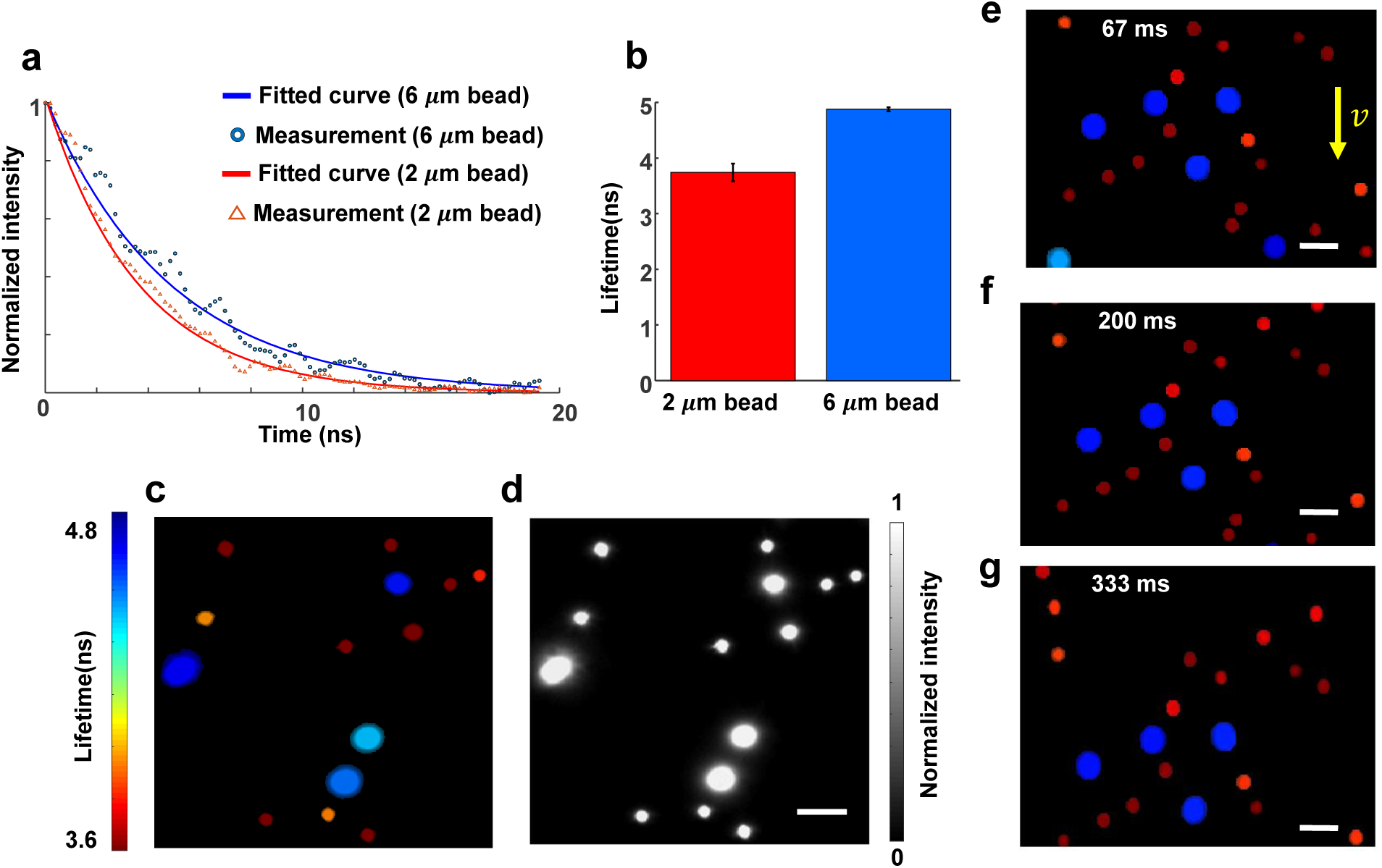
Lifetime imaging of fluorescent beads in flow. **a**. Reconstructed fluorescence decays of two types of fluorescent beads. **b**. Mean lifetimes. The standard errors of the mean are shown as error bars. **c**. Reconstructed snapshot lifetime image. **d**. Reference intensity image. **e-g**. Lifetime images of fluorescent beads in flow at representative temporal frames. Scale bar: 10 μm.

We further reconstructed the entire flow dynamics and show the movie and representative temporal frames in **Movie 2** and **Fig. 2e-g**, respectively. Because the lifetime image was acquired in a snapshot format, no motion blur is observed. Moreover, we calculated the average fluorescence lifetimes of these two types of beads. The results 4.9 ns and 3.7 ns match well with the ground truth.

### Lifetime unmixing of neural cytoskeletal proteins

Next, we applied compressed FLIM to cell imaging and demonstrated fluorescence lifetime unmixing of multiple fluorophores with highly overlapped emission spectra. Multi-target fluorescence labeling is commonly used to differentiate intracellular structures. Separation of multiple fluorophores can be accomplished by spectrally-resolved detection and multicolor analysis^36–38^ or time-resolved detection by FLIM^39-41^. The spectral method fails when the emission spectra of the fluorophores strongly overlap. By contrast, FLIM has a unique edge in this case provided a difference in fluorophores’ lifetimes.

We imaged two protein structures in the cytoskeleton of neurons and unmixed them based on the lifetime. We immunolabelled Vimentin and Tubulin in the cytoskeleton with Alexa Fluor 555 and Alexa Fluor 546, respectively. The emission spectra of two fluorophores highly overlap but their fluorescence lifetimes differ (1.3 ns *vs*. 4.1 ns). Within a single snapshot, we captured a high-resolution lifetime image of immunofluorescently-stained neurons (**Fig. 3a**). **Movie 3** records the time-lapse fluorescence decay process after a single pulse excitation, and two representative decay curves associated with Alexa Fluor 555 and 546 are shown in **Fig. 3b**. Next, we applied a regularized unmixing algorithm (Methods) to the lifetime data and separated Vimentin and Tubulin into two channels in **Fig. 3d** (green channel, Vimentin; blue channel, Tubulin). To acquire the ground-truth unmixing result, we operated our system in a slit-scanning mode and imaged the same field of view (Methods). The resultant unmixed result (**Fig. 3e**) matches well with compressed FLIM measurement. To further validate the distribution pattern of Vimentin and Tubulin in neuron cytoskeleton structures, we imaged the sample using a bench-mark confocal FLIM system (ISS Alba FCS), and **Fig. S1** presents the two proteins distribution. The neurons exhibit a similar protein distribution pattern as that inferred by compressed FLIM.

**Fig.3.**
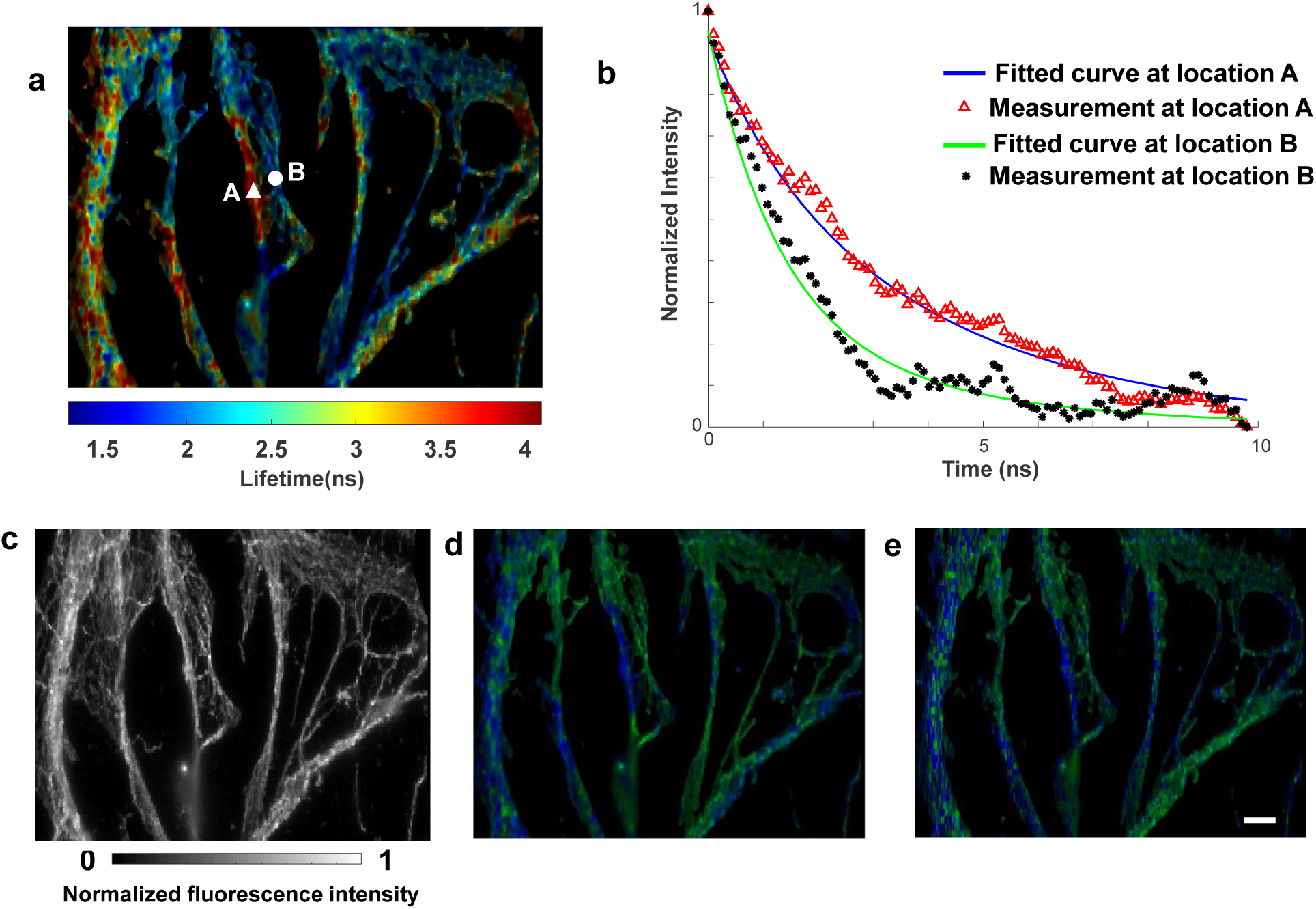
Lifetime imaging of neuronal cytoskeleton immunolabelled with two fluorophores. Alexa Fluor 555 immunolabels Vimentin. Alexa Fluor 546 immunolabels Tubulin. **a**. Reconstructed lifetime image of immunofluorescently-stained neurons. **b**. Reconstructed fluorescence decays at two fluorophore locations. **c**. Reference intensity image. **d**. Lifetime unmixed image. Green channel, Vimentin; Blue channel, Tubulin. **e**. Ground-truth lifetime unmixed image captured by line-scanning streak imaging. Scale bar: 10 µm.

### Imaging neural spikes in live cells

The complex functions of the brain depend on coordinated activity of multiple neurons and neuronal circuits. Therefore, visualizing the spatial and temporal patterns of neuronal activity in single neurons is essential to understand the operating principles of neural circuits. Recording neuronal activity using optical methods has been a long-standing quest for neuroscientists as it promises a noninvasive means to probe real-time dynamic neuronal function. Imaging neuronal calcium transients (somatic calcium concentration changes) with genetically encoded calcium indicators^42,43^, fluorescent calcium indicator stains^44^, and two-photon excitation methods using galvanometric^45^ and target-path scanners^46^ have been used to resolve suprathreshold spiking (electrical) activity.

However, calcium imaging is an indirect method of assessing neuronal activity, and spike number and firing rates using fluorescence recording is error prone, especially when used in cell populations that contain heterogeneous spike-evoked calcium signals. Thus, using functional calcium imaging to detect neuronal spiking in single-cell has been limited. Assessing neuronal function directly is invaluable for advancing our understanding of neurons and of the nervous system.

Genetically encoded voltage indicators (GEVIs) offer great promise for directly visualizing neural spike dynamics^47,48^. Compared with calcium imaging, GEVIs provide much faster kinetics that faithfully capture individual action potentials and sub-threshold voltage dynamics. Förster resonance energy transfer (FRET)-opsin fluorescent voltage sensors report neural spikes in brain tissue with superior detection fidelity compared with other GEVIs^49^.

As a molecular ruler, FRET involves the nonradiative transfer of excited state energy from a fluorophore, the donor, to another nearby absorbing but not necessarily fluorescent molecule, the acceptor. When FRET occurs, both the fluorescence intensity and lifetime of the donor decrease. So far, most fluorescence voltage measurements using FRET-opsin-based GEVIs report relative fluorescence intensity changes (Δ*F/F*) and fail to reveal the absolute membrane voltage because of illumination intensity variations, photobleaching, and background autofluorescence. By contrast, because FLIM is based on absolute lifetime measurement, it is insensitive to the environmental factors. Therefore, FLIM enables a more quantitative study with FRET-opsin-based GEVIs and provides a readout of the absolute voltage^50^.

To demonstrate compressed FLIM can be used to detect FRET, we first imaged two fluorescence dyes (donor, Alexa Fluor 546; acceptor, Alexa Fluor 647) in phosphate buffer solutions (PBS) with varied mixed concentration ratios. The emission spectrum of the donor overlaps considerably with the absorption spectrum of the acceptor, meeting the requirement for FRET. Acceptor bleed-through (ABT) contamination, *i*.*e*., the direct excitation and emission of the acceptor, is minimized by properly choosing the excitation wavelength and emission filter (Methods). We prepared three samples (**Fig. 4a**) with different concentration ratios (1:0, 1:1, 1:2) of donor and acceptor and imaged the fluorescence intensities and lifetimes using the reference time-integrated camera and compressed FLIM, respectively (**Fig. 4b-c**). As expected, fluorescence emission intensity of the donor gradually diminishes with more acceptor presence and stronger fluorescence quenching. Also, as revealed by compressed FLIM, there is a decrease in the donor’s fluorescence lifetime along with an increase in the acceptor’s concentration. Furthermore, we performed the ground-truth measurement by switching the system to the line-scanning mode (Methods). The lifetime results acquired by compressed FLIM match well with the ground truth (**Fig. 4d**).

**Fig.4.**
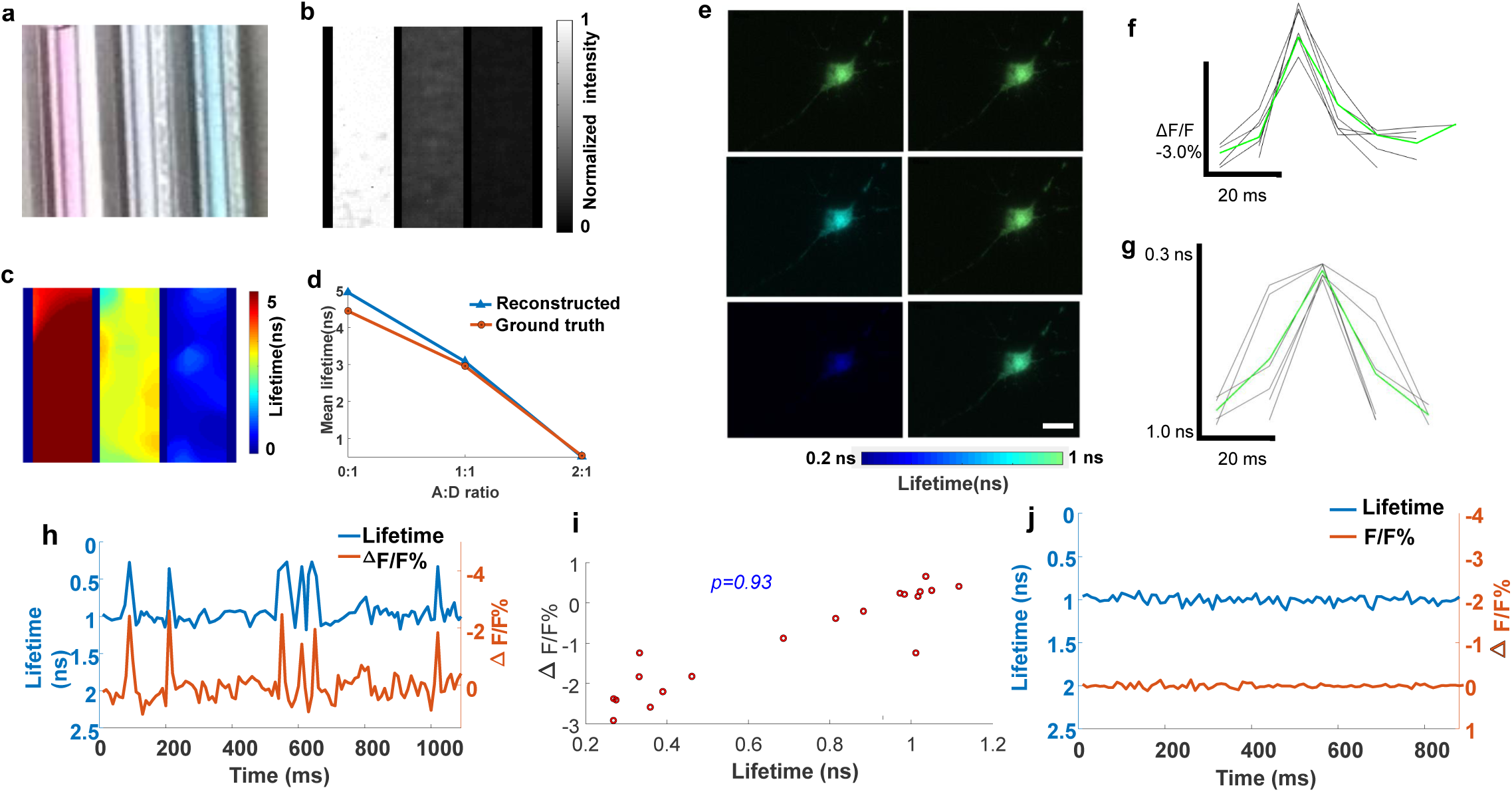
High-speed lifetime imaging of neural spiking in live neurons expressing MacQ-mOrange2. **a-d**. Lifetime imaging of FRET in phantoms. Acceptor Alexa 647 and donor Alexa 546 were mixed with varied concentration ratios (A:D ratio). **a**. Photograph of three mixed solutions with different A:D ratios (0:1, 1:1, 2:1). **b**. Reference intensity image. **c**. Reconstructed lifetime image. **d**. Comparison between measurement and ground truth. **e-j**. Lifetime imaging of FRET in live neurons. The FRET sensor MacQ-mOrange2 was expressed in a live neuron. **e**. Reconstructed lifetime images of the neuron at representative temporal frames upon potassium stimulation. **f-g**. Intensity and lifetime waveforms of neural spikes (black lines) and their mean (green line). **h**. Reconstructed time-lapse lifetime and intensity recording of neural spiking. The signals were averaged inside a cell. **i**. Scatter plot between lifetime and intensity of MacQ-mOrange2 measured at different times. The Pearson correlation coefficient is 0.93. **j**. Negative control. Scale bar: 10 µm.

Next, we evaluated compressed FLIM in imaging a FRET-opsin-based GEVI, MacQ-mOrange2^49^, to detect spiking in cultured neurons. During voltage depolarization, the optical readout is fluorescence quenching of the FRET donor mOrange2. We transfected neurons with plasmid DNA MacQ-mOrange2 and stimulated with high potassium treatment. We then used compressed FLIM to image the neural spikes.

To determine fluorescence lifetime and intensity traces for individual cells, we extracted the pixels that rank in the top 50% of the SNR values, defined as 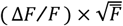, where (Δ*F/F*) is the voltage-dependent change in fluorescence intensity^49^, and 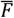 is a pixel’s mean baseline fluorescence intensity^49^. Average fluorescence lifetime and fluorescence intensity were calculated from these pixels in each frame. **Fig. 4h** shows the fluorescence intensity and lifetime traces of MacQ-mOrange2 sensor expressed in a cultured hippocampal neuron within 50 mM potassium environment imaged at 100 Hz. **Movie 4** records the lifetime dynamics due to neural pulsing. Representative snapshots at 50 ms, 60 ms, 70 ms, 80 ms, 90 ms and 100 ms shown in **Fig. 4e** indicate occurrence of lifetime oscillations. The observed spiking irregularity may attribute to ion channel stochasticity^51^, recurrent activity in the neuronal network^52^, or modulation of neuronal excitability^53^. The average relative fluorescence intensity change (Δ*F/F*) and absolute lifetime change (Δτ) in response to one spiking event are −2.9% and −0.7 ns, respectively. **Fig. 4f-g** present the experimentally determined fluorescence intensity and lifetime waveforms of single action potentials from MacQ-mOrange2 (black trace) and their mean (green trace, average over n=6 spikes). To further study the correlation between fluorescence intensities and lifetimes, we scatter plotted their relationship in **Fig. 4i**. The calculated Pearson coefficient is 0.93, indicating a high correlation between measured fluorescence intensities and lifetimes. Finally, to provide a negative control, we imaged MacQ-mOrange2 within a subthreshold non-activated 20 μM potassium stimulation (**Fig. 4j**). Both the fluorescence intensities and lifetimes were stable during the entire time trace, and no oscillations were observed. Because advancement in imaging speed is crucial for resolving the dynamics of neural activity at the single cell and across neural networks, the results presented here demonstrated the utility of compressed FLIM in neuroimaging.

## Discussion

Compared with conventional scanning-based FLIM imagers, compressed FLIM features three predominant advantages. First, based on a compressed-sensing architecture, compressed FLIM can produce high-resolution 2D fluorescence lifetime maps at 100 fps, allowing quantitative capture of transient biological dynamics. The gain in the imaging speed is attributed to the compressibility of a fluorescence scene in a specific domain. To show the dependence of reconstructed image quality on the compression ratio (CR) of a fluorescence event, we calculated the CR when imaging a biological cell stained with a typical fluorophore with a lifetime of 4 ns. The observation time window on the streak camera was set as 20 ns. Here we define CR as the ratio of the total number of voxels (*N*_*x*_ × *N*_*y*_ × *N*_*z*_; *N*_*x*_, *N*_*y*_, *N*_*t*_, samplings along spatial axes *x, y* and temporal axis *t*, respectively) in the reconstructed event datacube to the total number of pixels (*N*_*x′*_ × *N*_*y′*_; *N*_*x′*_, *N*_*y′*_, samplings along spatial axes *x′, y′*in the camera coordinate, respectively) in the raw image:

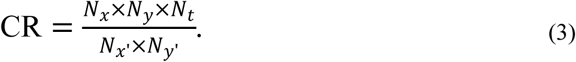

In compressed FLIM, the spatial information *y* and temporal information *t* occupy the same axis *y′*in the raw image. Therefore, their sum, *N*_*y*_ + *N*_*t*_ − 1, cannot exceed the total number of camera pixels along *y′* axis. Here we consider the equality case, *N*_*y′*_ = *N*_*y*_ + *N*_*t*_ − 1. Also, for simplicity, we assume the two complementary image channels fully occupy the entire *x′* axis on the camera, *i*.*e*., *N*_*x′*_ = 2*N*_*x*_. We then rewrite CR as:

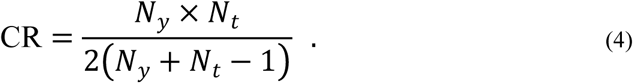

To evaluate the relation between CR and the reconstructed image quality, we used the peak signal-to-noise (PSNR) as the metric. For a given output image format (*N*_*y*_ = 200 pixels), we varied the CR by increasing *N*_*t*_ and calculated the corresponding PSNRs (**Fig. S3a**). As expected, increasing CR unfavorably reduces the reconstructed image quality. Therefore, in compressed FLIM, the sequence depth of the reconstructed event datacube must be balanced for image quality.

Second, because of using a complementary encoding scheme, compressed FLIM possesses a 100% light throughput (ignoring reflection losses from the optical elements). The light throughput advantage can be characterized by the snapshot advantage factor, which is defined as the portion of datacube voxels that are continuously visible to the instrument^54^. When measuring a high-resolution image in the presented format (500 × 400 pixels), we gain a factor of 2×10^5^ in light throughput compared with that in its point-scanning-based counterpart. Such a throughput advantage makes compressed FLIM particularly suitable for low light imaging applications. To study the dependence of the reconstructed image quality on the number of photons received at a pixel, we simulated the imaging performance of compressed FLIM for a shot-noise-limited system. Provided that the pixel with the maximum count in the time-integrated image channel collects ***M*** photons, the corresponding shot noise is 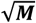 photons. We added photon noises to all pixels both in the temporally-sheared and integrated images and performed the image reconstruction with various ***M***s. **Fig. S3b** shows that a larger ***M*** (i.e., more photons) leads to a higher PSNR. For high-quality image reconstruction (PSNR≥20 dB), ***M*** must be greater than 80 photons.

Lastly, operating in a snapshot format, compressed FLIM eliminates the motion artifacts and enables fast recording of stochastic biological events. We quantitatively computed the maximum blur-free motion allowed by our system. Assuming we image a typical fluorophore using a 1.4 NA objective lens and a 20 ns observation time window on the streak camera, the maximum blur-free movement during a single-shot acquisition equals the system’s spatial resolution (∼0.2 *μ*m). The correspondent speed limit is 10 μm/μs. A lower NA objective lens or a shorter observation window will increase this threshold, however, at the expense of a reduced resolution and temporal sequence depth. The ability to capture rapid motion at the microscopic scale will be valuable to studying fast cellular events, such as protein folding^55^ or ligand binding^56^. Moreover, compressed FLIM employs widefield illumination to excite the sample, presenting a condition that is favorable for live cell imaging. Cells exposed to radiant energy may experience physiological damages because of heating and/or the generation of reactive oxygen species (ROS) during extended fluorescence microscopy^57^. Because phototoxicity has a nonlinear relation with illumination radiance^58^, widefield compressed FLIM prevails in preserving cellular viability compared with its scanning-based counterpart.

In compressed FLIM, the spatial and temporal information are multiplexed and measured by the same detector array. The system therefore faces two constrains. First, there is a trade-off between lifetime estimation accuracy and illumination intensity. Given ample photons, increasing the number of temporal samplings *N*_*t*_ will improve the lifetime estimation accuracy approximately in the manner of 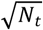. Using a faster temporal shearing velocity in the streak camera will disperse the fluorescence signals to more detector array rows at the expense of a reduced signal-to-noise ratio. To maintain the photon budget at each temporal sampling bin, we must accordingly increase the illumination intensity at the sample, a fact which may introduce photobleaching and shorten the overall observation time. The second trade-off exists between *y*-axis spatial sampling and *t*-axis temporal sampling. For a given format detector array, without introducing temporal shearing, the final raw image occupies *N*_*y*_ pixel rows. With temporal shearing, the image is sheared to *N*_*y*_ + *N*_*t*_ − 1 pixel rows with the constraint *N*_*y*_ + *N*_*t*_ − 1≤*N*_*y*’_. Here *N*_*y′*_ is the total number of detector rows. Increasing the image size *N*_*y*_ will decrease the *N*_*t*_ and thereby the lifetime estimation accuracy. Therefore, the image FOV and lifetime estimation accuracy must be balanced for a given application.

In summary, we have developed a high-speed, high-resolution fluorescent lifetime imaging method, compressed FLIM, and demonstrated its utility in imaging dynamics. Capable of capturing a 2D lifetime image within a snapshot, we expect compressed FLIM would have broad applications in blur-free observation of transient biological events, enabling new avenues of both basic and translational biomedical research.

## Methods

### Forward model

We describe compressed FLIM’s image formation process using the forward model^5^. Compressed FLIM generates three projection channels: a time-unsheared channel, and two time-sheared channels with complementary encoding. Upon laser illumination, the fluorescence decay scene is first imaged by the microscope to the conjugated plane and *I*(*x,y,t*) denotes the intensity distribution. A beam splitter then divides the conjugated dynamic scene into two beams. The reflected beam is directly captured by a reference CMOS camera, generating a time-integrated image. The optical energy, *E*_*I*_(*m,n*), measured at pixel *m, n*, on the reference CMOS camera is:

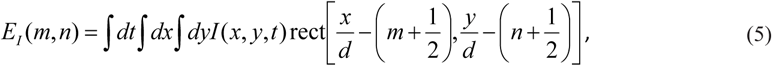

where *d* is the pixel size of the reference camera.

The transmitted beam is then relayed to an intermediate plane (DMD plane) by an optical imaging system. Assuming unit magnification and ideal optical imaging, the intensity distribution of the resultant intermediate image is identical to that of the original scene. The intermediate image is then spatially encoded by a pair of complementary pseudorandom binary patterns displayed at the DMD plane. The two reflected spatially-encoded images have the following intensity distribution:

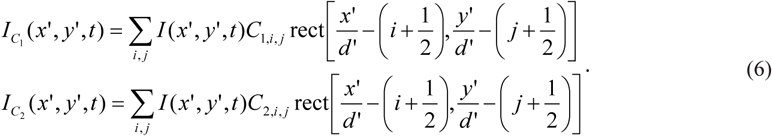

Here, *C*_*1,i,j*_ and *C*_*2,i,j*_ are elements of the matrix representing the complementary patterns with *C*_*1,i,j*_ + *C*_*2,i,j*_ =1, *i, j* are matrix element indices, and *d’* is the binned DMD pixel size. For each dimension, the rectangular function (rect) is defined as:

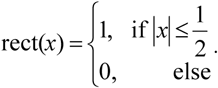

The two reflected light beams masked with complimentary patterns are then passed to the entrance port of a streak camera. By applying a voltage ramp, the streak camera acts as a shearing operator along the vertical axis (*y′’* axis in **Fig. 1**) on the input image. Assuming ideal optics with unit magnification again, the sheared images can be expressed as

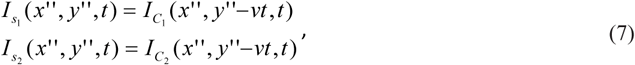

where *v* is the shearing velocity of the streak camera.

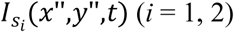 is then spatially integrated over each camera pixel and temporally integrated over the exposure time. The optical energy, 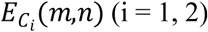, measured at pixel *m, n*, is:

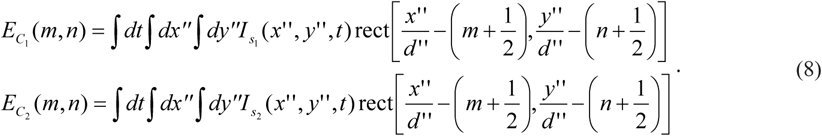

Here, *d’’* is the camera pixel size. Accordingly, we can voxelize the input scene, *I*(*x,y,t*), into *I*_*i,j,k*_ as follows:

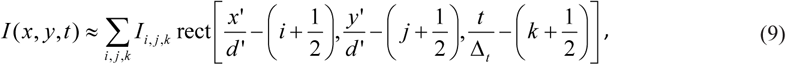

where *Δ*_*t*_ *= d’’/v*. If the pattern elements are mapped 1:1 to the camera pixels (that is, *d’=d’’*) and perfectly registered, and the reference CMOS camera and the internal CMOS camera of the streak camera have the same pixel size (that is, *d =d’’*), combining equations (5)-(9) yields:

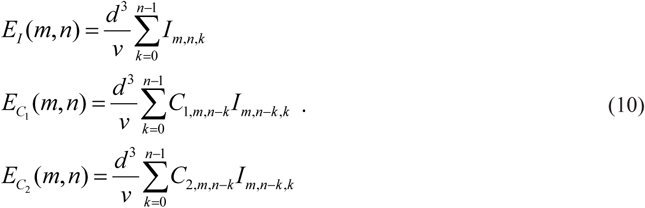

Here *C*_*i,m,n-k*_ *I*_*m,n-k,k*_ (i = 1, 2) represents the complimentary-coded, sheared scene, and the inverse problem of equation (10) can be solved using existing compressed-sensing algorithms^35,59,60^.

### Compressed FLIM image reconstruction algorithm

Given prior knowledge of the binary pattern, to estimate the original scene from the compressed FLIM measurement, we need to solve the inverse problem of equation (10). Because of the sparsity in the original scene, the image reconstruction can be realized by solving the following optimization problem

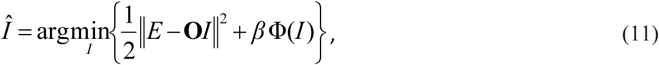

where ***O*** is the linear operator, Φ(*I*) is the regularization function and *β* is the weighing factor between the fidelity and sparsity. To further impose space and intensity constraint, we construct the new constrained solver:

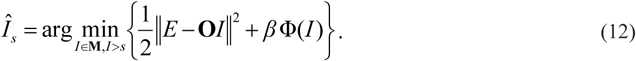

Here, **M** is a set of possible solutions confined by a spatial mask extracted from the reference intensity image and defines the zone of action in the reconstruction. This spatial constraint improves the image resolution and accelerates the reconstruction. *s* is the low intensity threshold constraint to reduce the low-intensity artifacts in the data cube. In compressed FLIM image reconstruction, we adopt an algorithm called two-step iterative shrinkage/thresholding(TwIST)^35^, with Φ(*I*) in the form of total variation (TV):

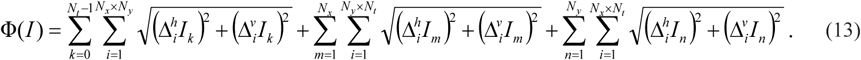

Here we assume that the discretized form of *I* has dimensions *N*_*x*_ × *N*_*y*_ × *N*_*t*_ (*N*_*x*_, *N*_*y*_ and *N*_*t*_ are respectively the numbers of voxels along *x, y* and *t*), and *m, n, k* are three indices. *I*_*m*_, *I*_*n*_, *I*_*k*_ denote the 2D lattices along the dimensions *m, n, k*, respectively. 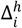 and 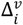 are horizontal and vertical first-order local difference operators on a 2D lattice. After the reconstruction, non-linear least squares exponential fitting is applied to data cube along the temporal dimension at each spatial sampling point to extract the lifemap. As shown in **Fig. 2a**, compressed FLIM reconstructs a fluorescence decay process at one sampling point which matches well with the ground truth.

### Lifetime-based fluorophore un-mixing algorithm

For simplicity, here we consider only fluorophores with single-exponential decays. Provided that the sample consists of *n* mixed fluorophores with lifetimes ***τ*** *=* (*τ1, …, τn*) and concentration ***x*** = (*x1, …, xn*), upon a delta pulse excitation, the discretized time-lapse fluorescence decay is:

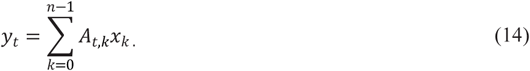

Here, *k* is the fluorophore index, *A*_*t,k*_ is an element of the fluorescence decay component matrix ***A***, and *A*_*t,k*_ = exp (−*t*/τ_*k*_). The inverse problem of equation (14) is a least squares problem with constraints. We choose 2-norm penalty and form the solvent for *ℓ*_*2*_-regularized least squares problem:

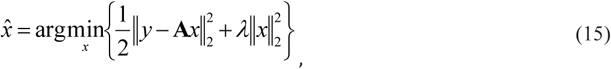

where the first term 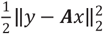 represents the measurement fidelity, and the regularization term penalizes large norm of *x*. The regularization parameter *λ* adjusts the weight ratio between fidelity and 2-norm penalty.

In our experiment, to construct fluorescence decay component matrix ***A***, we first directly imaged Alexa Fluorophore 555 and Alexa Fluorophore 546 in solution and captured their time-lapse fluorescence decay. Then we computed their lifetimes by fitting the asymptotic portion of the decay data with single exponential curves. Finally, we applied the regularized unmixing algorithm to the lifetime data and separated the fluorophores into two channels.

### GPU assisted real-time reconstruction using computer cluster

Based on the iterative construction, compressed FLIM is computationally extensive. For example, to reconstruct a 500 × 400 × 617 (*x, y, t*) event datacube and compute a single lifetime image, it takes tens of minutes on a single PC. The time of constructing a dynamic lifetime movie is prohibitive. To accelerate this process, we (1) implemented the reconstruction algorithm using a parallel programming framework on two NVIDIA Tesla K40 GPUs and (2) performed all reconstructions simultaneously on a computer cluster (Illinois Campus Cluster). The synergistic effort significantly improved the reconstruction speed and reduced the movie reconstruction time to seconds. **Table 1** illustrates the improvement in reconstruction time when the computation is performed on a single PC vs. the GPU-assisted computer cluster.

**Table 1.** Reconstruction time comparison when the computation is performed on a single PC vs. a GPU-assisted computer cluster

|  | Single PC | GPU-assisted computer cluster |
| --- | --- | --- |
| Single lifetime image | 155s | 2.7s |
| Lifetime movie (100 frames) | 4.3 hours | 2.7s |

### Compressed FLIM: hardware

In the compressed FLIM system, we used an epi-fluorescence microscope (Olympus IX83) as the front-end optics (**Fig. S4a**). We excited the sample using a 515 nm picosecond pulse laser (Genki-XPC, NKT Photonics) and separated the fluorescence from excitation using a combination of a 532 nm dichroic mirror (ZT532rdc, Chroma) and a 590/50 nm band-pass emission filter (ET590/50m, Chroma). Upon excitation, an intermediate fluorescence image was formed at the side image port of the microscope. A beam splitter (BSX16, Thorlabs) transmitted 10% of light to a temporal-integration camera (CS2100M-USB, Thorlabs) and reflected 90% of light to the temporal-shearing channels. The reflected image was then relayed to a DMD (DLP LightCrafter 6500, Texas Instruments) through a 4f system consisting of a tube lens (AC508-100-A, Thorlabs) and a stereoscopic objective (MV PLAPO 2XC, Olympus; numerical aperture, 0.50). At the DMD, we displayed a random, binary pattern to encode the image. The reflected light from both the “on” mirrors (tilted +12*°* with respect to the norm) and “off” mirrors (tilted −12° with respect to the norm) were collected by the same stereoscopic objective, forming two complementary channel images at the entrance port of a streak camera (C13410-01A, Hamamatsu) (**Fig. S4b**). The streak camera deflected the image along the vertical axis depending on the time-of-arrival of incident photons. The resultant spatially-encoded, temporally-sheared images were acquired by an internal CMOS camera (ORCA-Flash 4.0 V3, C13440-20CU, Hamamatsu) with a sensor size of 1344(H) × 1016(V) pixels (1 × 1 binning; pixel size *d* = 6.5 µm). We synchronized the data acquisition of cameras using a digital delay generator (DG645, Stanford Research Systems).

### Filter selection for FRET-FLIM imaging

To assure only fluorescence emission from the donor was collected during FRET-FLIM imaging, we chose a filter set (a 515 nm excitation filter and a 590/50 nm emission filter) to separate excitation and fluorescence emission. This filter combination suppressed the direct excitation of the acceptor to <3% and minimized the collection of the acceptor’s fluorescence (**Fig. S5**), thereby eliminating the acceptor bleed-through (ABT) contamination.

### Spatial registration among three imaging channels

Because compressed FLIM imaged a scene in three channels (one temporal-integrated channel and two complementary temporal-shearing channels), the resultant images must be spatially registered. We calibrated the system using a point-scanning-based method. We placed an illuminated pinhole at the microscope’s sample stage and scanned it across the FOV. At each point-scanning position, we operated the streak camera in the “focus” mode (i.e., without temporal shearing) and captured two impulse response images with all DMD’s pixels turned “on” and “off”, respectively. Meanwhile, the reference CMOS camera captured another impulse response image in the temporal-integration channel. We then constructed a lookup table by extracting the pinhole locations in these three impulse response channel images. This lookup table was later used to register the three channel images 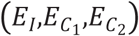 for concatenated image reconstruction.

### Acquisition of encoding matrices C_1_ and C_2_

To acquire the encoding matrices **C**_1_ and **C**_2_, we imaged a uniform scene and operated the streak camera in the “focus” mode. The streak camera directly captured the encoding patterns without temporal shearing. Additionally, we captured two background images with all DMD’s pixels turned “on” and “off”, respectively. To correct for the non-uniformity of illumination, we then divided the coded pattern images by the corresponding background images pixelwise.

### Slit-scanning streak camera imaging

To form a ground-truth lifetime image, we employed the DMD as a line scanner and scanned the sample along the direction perpendicular to the streak camera entrance slit. We turned on the DMD’s (binned) mirror rows sequentially and imaged the temporally-sheared line image in the correspondent imaging channel. Given no spatiotemporal mixing along the vertical axis, the fluorescence decay data along this line direction could be directly extracted from the streak image. Next, we computed the fluorescence lifetimes by fitting the decay data to single exponential curves. The resultant line lifetime images were stacked to form a 2D representation.

### Confocal FLIM imaging

For confocal FLIM imaging, we used a bench-mark commercial system (ISS Alba FCS). The sample was excited by a Ti:Sapphire laser, and fluorescence was collected by a Nikon Eclipse Ti inverted microscope. The time-lapse fluorescence decay was measured by a time-correlated single photon counting unit. To form a 2D lifetime image, the system raster scanned the sample. The typical time to capture a 2D lifetime image (256 × 256 pixels) was ∼ 60 s.

### Fluorescence beads

We used a mixture of 6 μm and 2 μm diameter fluorescence beads (C16509, Thermo Fisher; F8825 Thermo Fisher) in our experiment. To prepare the mixed beads solution, we first diluted the 6 μm and 2 μm diameter bead suspensions. After sonicating the two fluorescence bead suspensions, we pipetted 10 uL of the 6 μm bead suspension (∼1.7 × 10^7^ beads/mL) and 2 μm bead suspension (∼4.5 × 10^9^ beads/mL) into 1mL and 10mL phosphate buffer solutions (PBS), respectively. Next, we mixed 100 uL diluted 2 μm bead solution with 1 ml diluted 6 μm bead solution. The final mixed beads solution contained approximately 1.5 × 10^5^ 6 μm beads/mL and 4.1 × 10^5^ 2 μm beads/mL.

### FRET phantom

We used Alexa Fluor 546 (A-11003, Thermo Fisher) as the donor and Alexa Fluor 647 (A-21235, Thermo Fisher) as the acceptor. We prepared the acceptor solutions with three different concentrations (0 mg/mL, 1 mg/mL, 2 mg/mL) and mixed them with the same donor solution(1 mg/mL). We then injected them into three glass tubes (14705-010, VWR) for imaging.

### Primary cell culture

Primary hippocampal neurons were cultured from dissected hippocampi of Sprague-Dawley rat embryos. Hippocampal neurons were then plated on 29mm glass bottom petri dishes that are pre-coated with poly-D-lysine (0.1 mg/ml; Sigma-Aldrich). To help the attachment of neurons (300 cells/mm^2^) on to the glass bottom dish, neurons were initially incubated with a plating medium containing 86.55% Minimum Essential Medium Eagle’s with Earle’s Balanced Salt Solution (MEM Eagle’s with Earle’s BSS, Lonza), 10% Fetal Bovine Serum (re-filtered, heat inactivated FBS; ThermoFisher), 0.45% of 20% (wt./vol.) glucose, 1**×** 100 mM sodium pyruvate (100**×**; Sigma-Aldrich), 1**×** 200 mM glutamine (100**×**; Sigma-Aldrich), and 1**×** penicillin/streptomycin (100**×**; Sigma-Aldrich). After three hours of incubation in the incubator (37 ° C and 5% CO_2_), the plating media was aspirated and replaced with maintenance media containing Neurobasal growth medium supplemented with B-27 (Invitrogen), 1% 200 mM glutamine (Invitrogen) and 1% penicillin/streptomycin (Invitrogen) at 37 ° C, in the presence of 5% CO_2_. Half the media was aspirated once a week and replaced with fresh maintenance media. The hippocampal neurons were grown for 10 days before imaging.

### Immunofluorescently stained neurons

We immunolabelled the Vimentin (MA5-11883, Thermo Fisher) with Alexa Fluor 555 (A-21422, Thermo Fisher) and Tubulin (PA5-16891, Thermo Fisher) with Alexa Fluor 546 (A-11010, Thermo Fisher) in the neurons.

### Transfect the neurons to express MacQ-mOrange2

We used Mac mutant plasmid DNA Mac-mOrange2 (#48761, Addgene) to transfect hippocampal neurons. In DNA collection, as soon as we received the agar stab, we grew bacteria in Luria-Bertani (LB) broth with ampicillin in 1:1000 dilution for overnight in 37°C. Standard Miniprep (Qiagen) protocol was performed in order to collect DNA. DNA concentration was measured by Nanodrop 2000c (ThermoFisher). In neuron transfection, lipofectamine 2000 (Invitrogen) was used as transfecting reagent. In an Eppendorf tube, we stored 1mL of the conditioned culture media from 29mm petri plates neuron culture with 1mL of fresh media. We prepared two separate Eppendorf tubes and add 100uL of Neurobasal medium to each tube. For one tube, 3ug of DNA was added while 4uL of lipofectamine 2000 was added to the other tube. After five minutes, two tubes were mixed together and incubated in room temperature for 20 minutes. This mixture was added to neuron culture in dish for 4 hours in the incubator (37°C and 5% CO_2_). We took out the media containing lipofectamine 2000 reagent and added the stored conditioned and fresh culture media to neuron culture in dish. Hippocampal neurons were imaged 40 hours after transfection.

### Potassium stimulation and imaging

We used high potassium (50mM) treatment to stimulate neuron spike. The extracellular solution for cultured neurons (150 mM NaCl, 4 mM KCl, 10 mM glucose, 10 mM HEPES, 2 mM CaCl_2_ and 2 mM MgCl_2_) was adjusted to reach the desired final K^+^ concentration (50mM) and maintain physiological osmolality at the same time. At each stimulation, we removed the media in the plate and pipetted high potassium extracellular solution at the same time.

## Data availability

The data that support the plots within this paper and other findings of this study are available from the corresponding author upon reasonable request.

## Code availability

The code that support the findings of this study are available from the corresponding author upon reasonable request.

## Acknowledgement

This work was supported partially by National Institutes of Health (R01EY029397, R35GM128761); National Science Foundation (1652150).

## Author information

## Integrated supplementary information

**Fig.S1.**
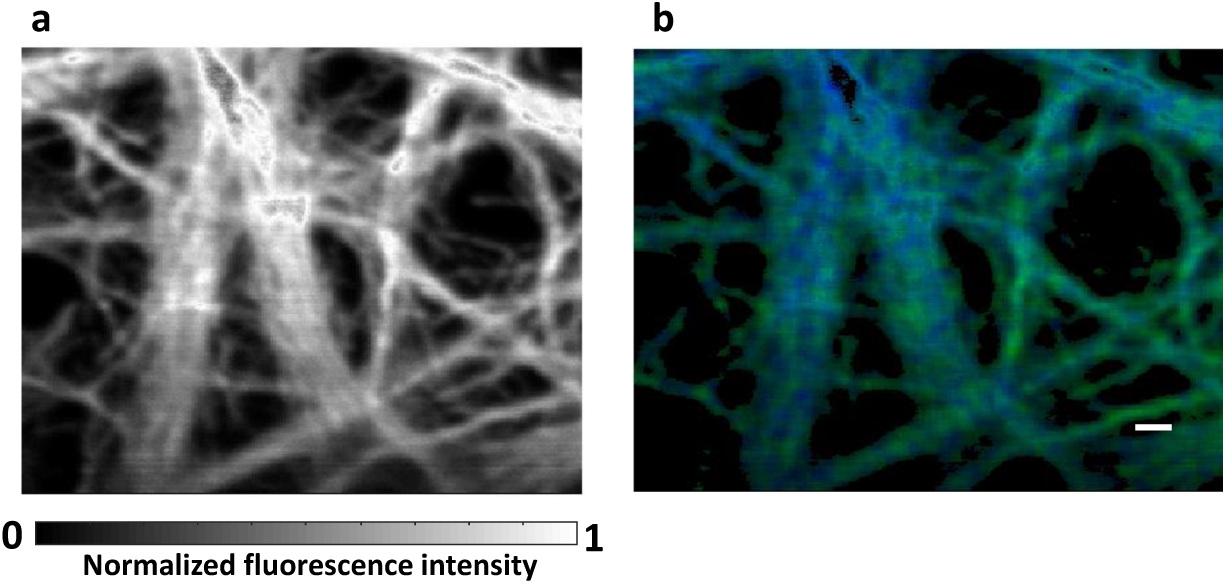
Confocal FLIM of cytoskeleton immunolabelled with two fluorophores. Alexa 555 immunolabels Vimentin. Alexa 546 immunolabels Tubulin. **a**. Intensity image. **b**. Lifetime unmixed image. Green channel, Vimentin; Blue channel, Tubulin. Scale bar: 10 µm.

**Fig.S2.**
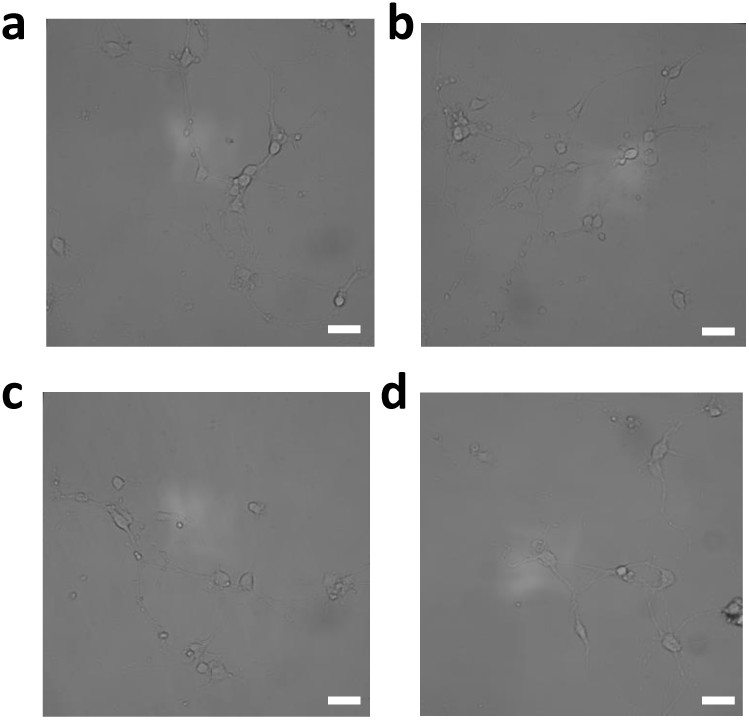
Brightfield images of cultured neurons expressing MacQ-mOrange2 under ×20 0.7 NA objective. Scale bar: 20 µm.

**Fig.S3.**
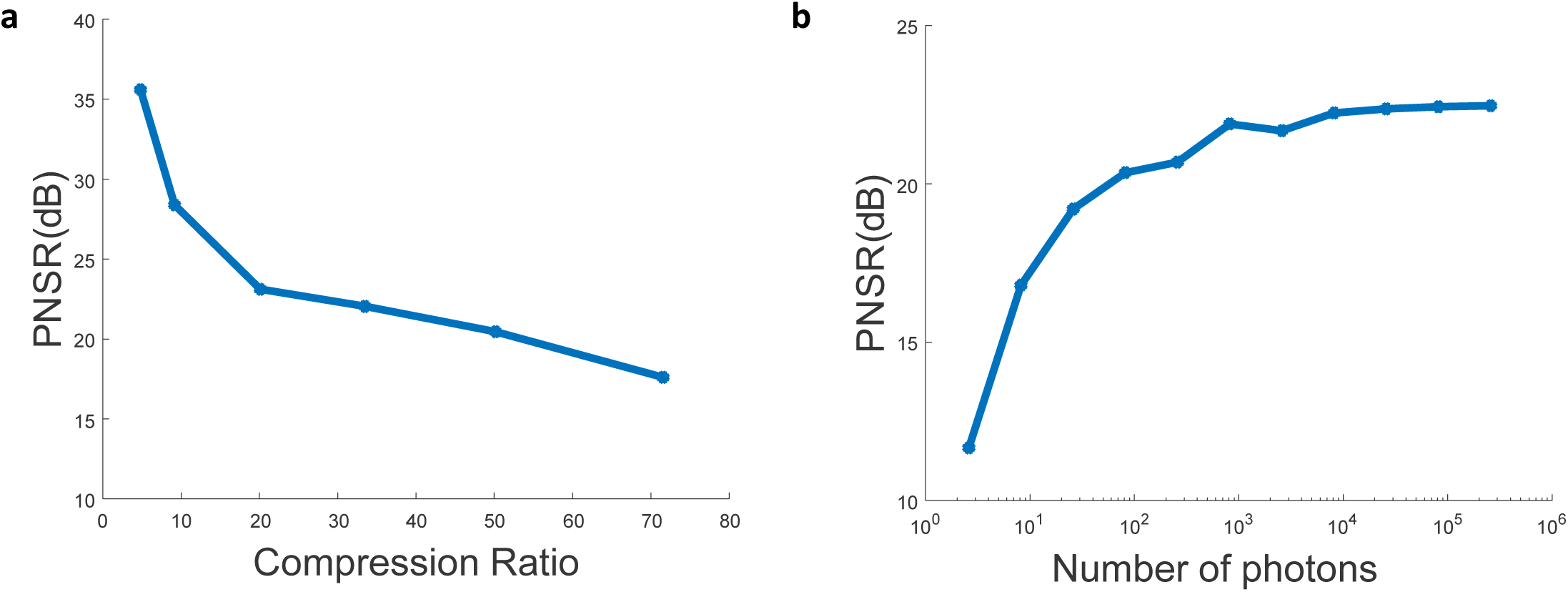
Dependence of the peak signal-to-noise (PSNR) on the compression ratio and the number of photons received at each pixel. **a**. PSNR vs. compression ratio. **b**. PSNR vs. number of photons received at a pixel.

**Fig.S4.**
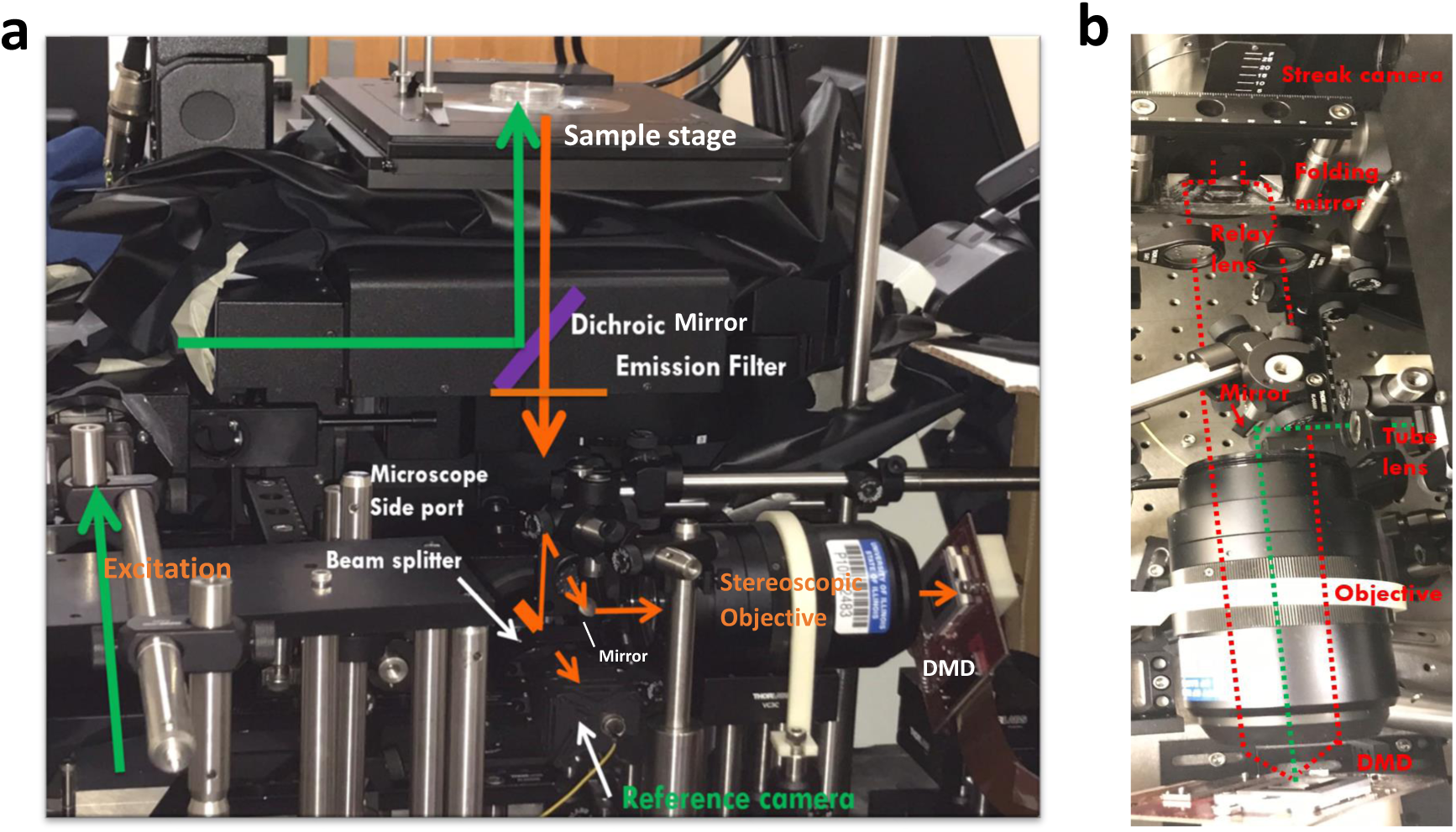
Photograph of the compressed FLIM system. **a**. Side view. **b**. Top view.

**Fig.S5.**
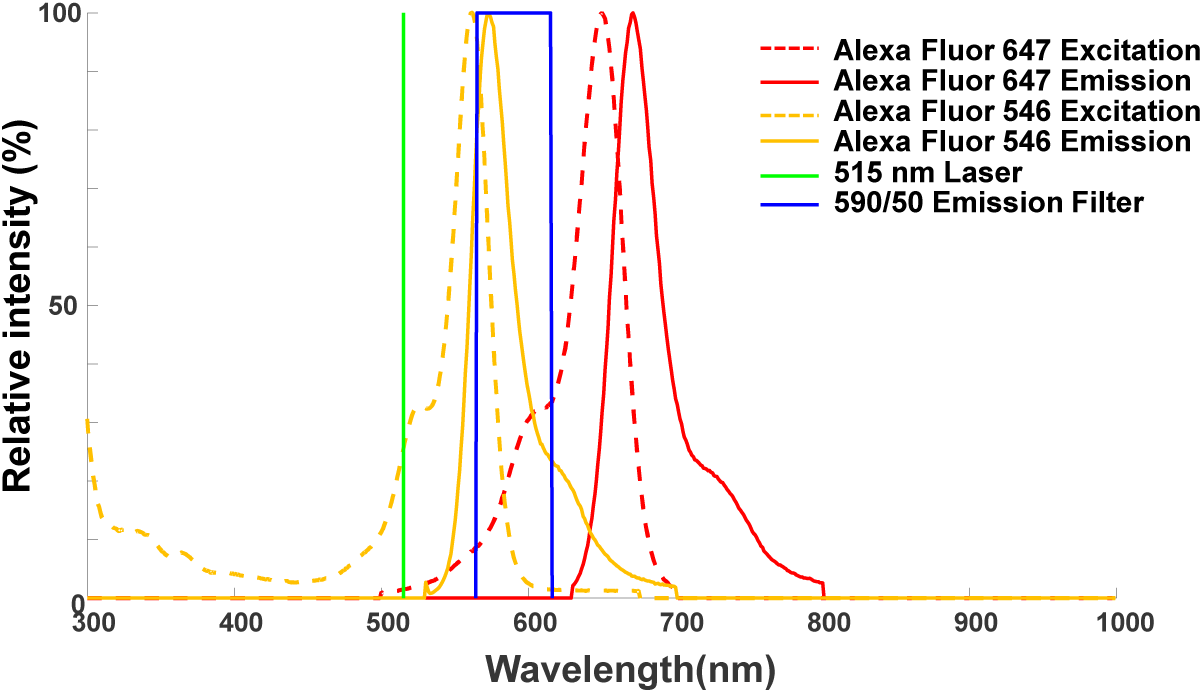
Filter selection for FRET-FLIM imaging.

**Fig.S6.**
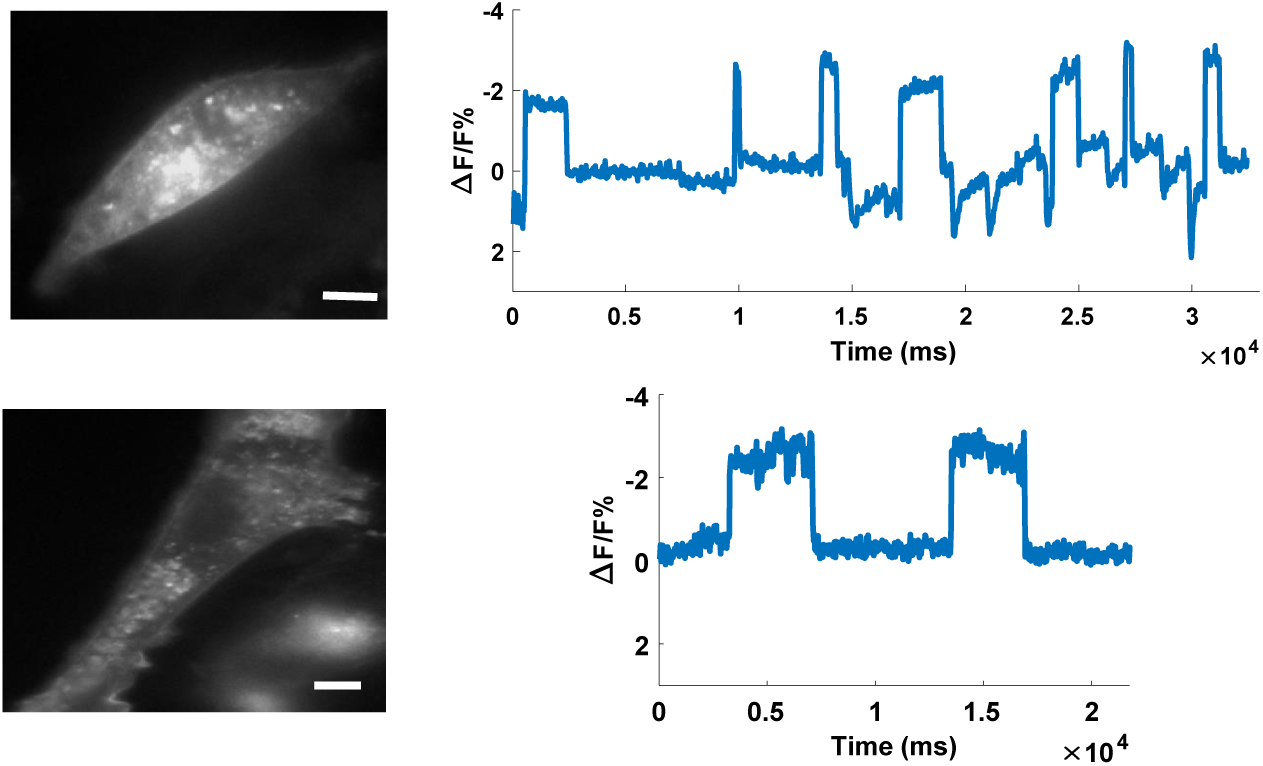
Intensity images and traces of two cultured rat heart cardiomyocytes expressing MacQ-mOrange2 stimulated by acetylcholine. Scale bar: 10 µm.

**Fig.S7.**
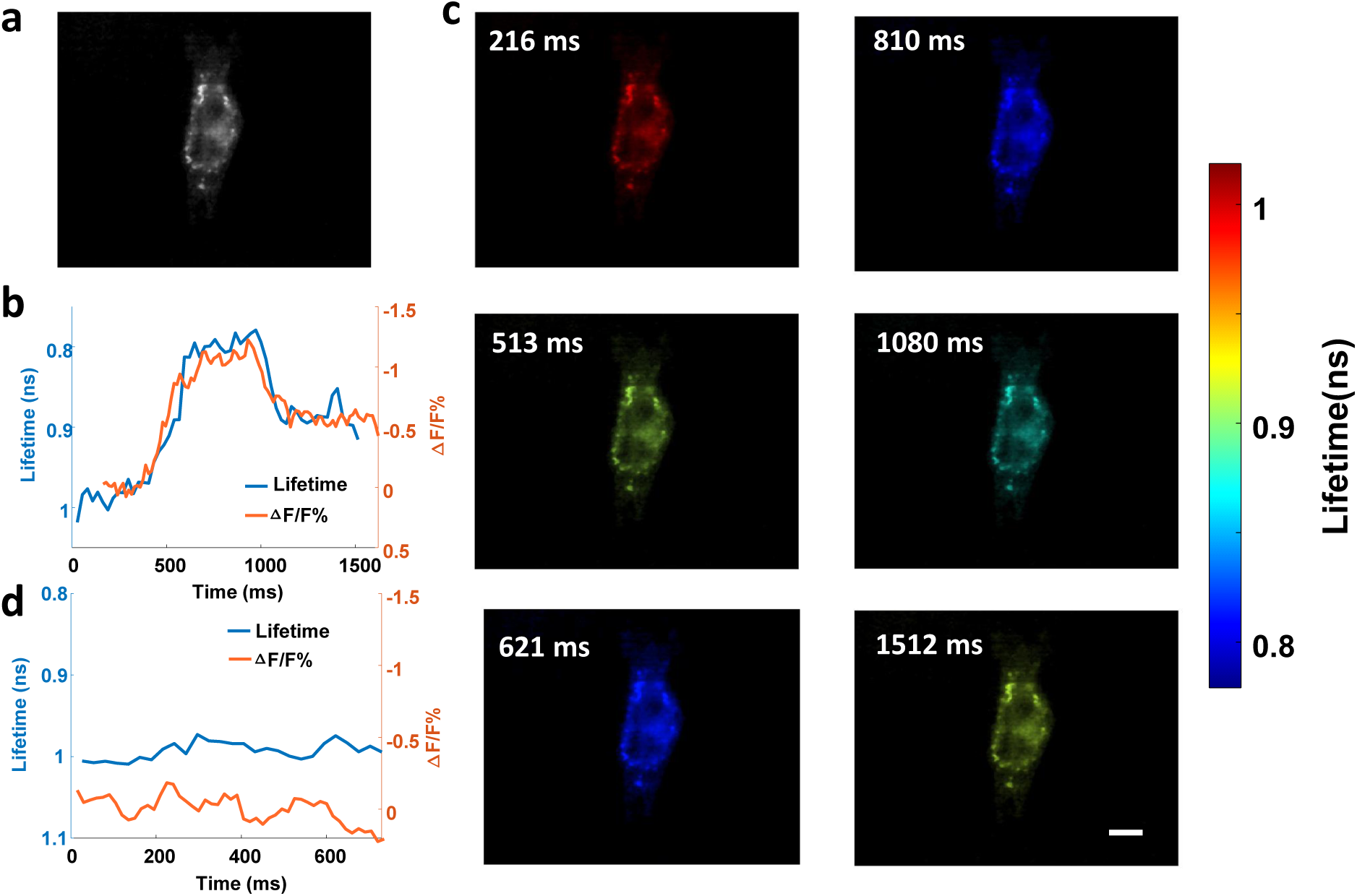
High-speed lifetime imaging of cardiomyocyte beating in live rat heart cardiomyocytes expressing MacQ-mOrange2. **a**. Intensity image. **b**. Reconstructed time-lapse lifetime and intensity recording of cardiomyocyte beating upon acetylcholine stimulation. **c**. Reconstructed lifetime images of the cardiomyocyte at representative temporal frames. **d**. Negative control. Scale bar: 10 µm.

**Fig.S8.**
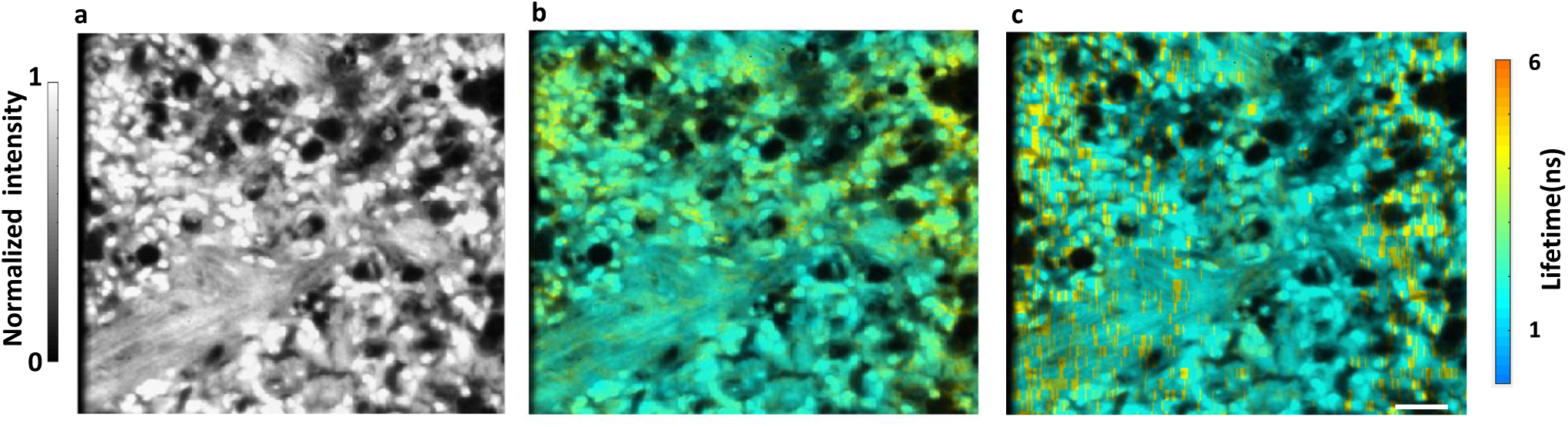
Lifetime imaging of a H&E slide. **a**. Intensity image. **b**. Reconstructed lifetime image. **c**. Ground truth. Scale bar: 20 μm.

**Fig.S9.**
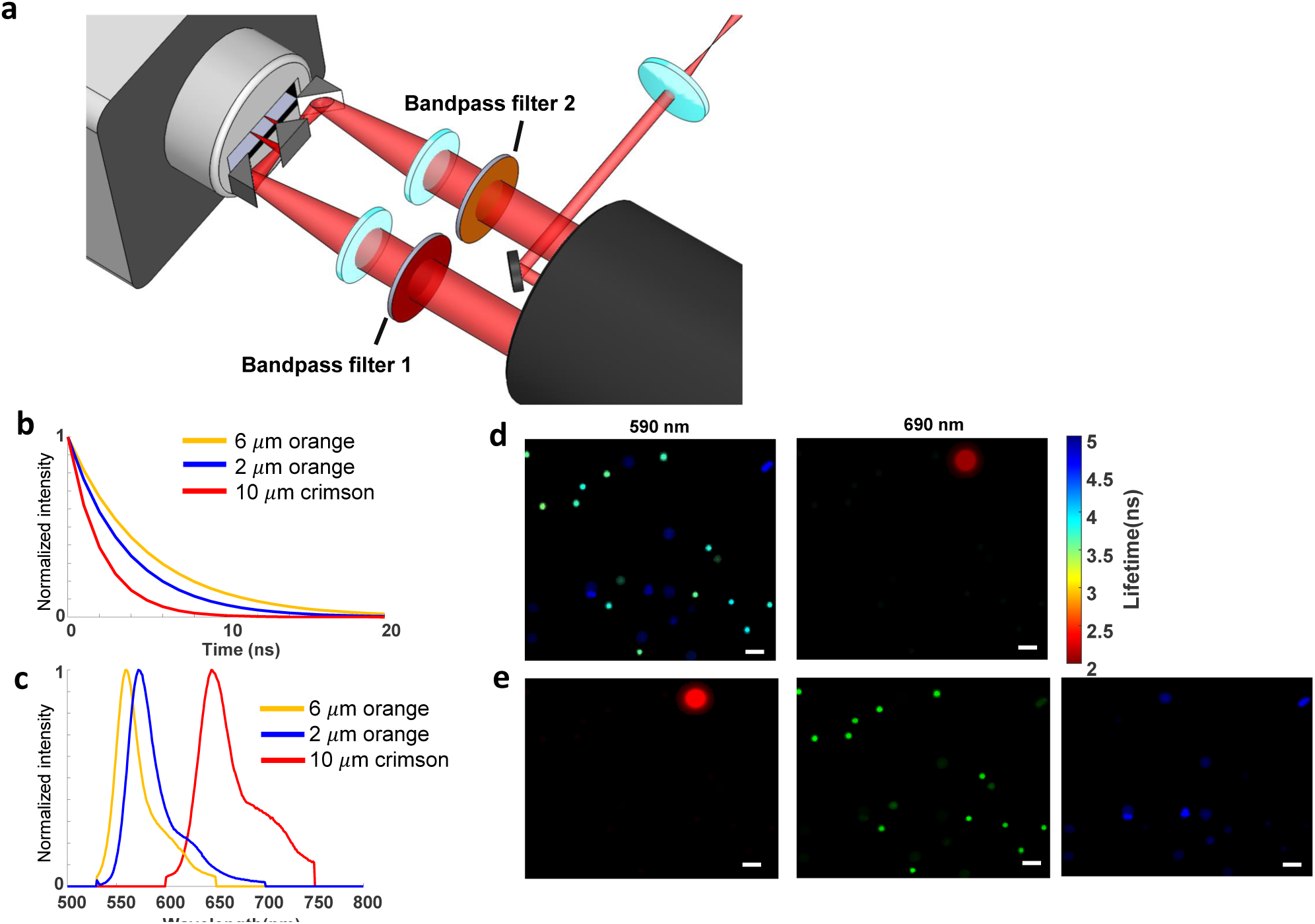
Compressed sFLIM for multi-target fluorescence imaging. **a**. Schematic. **b**. Time-lapse fluorescence decays of fluorophores contained in three types of fluorescence beads. **c**. Corresponding emission spectra of orange and crimson fluorophores. **d**. Reconstructed lifetime images at 590 nm and 690 nm. **e**. Spectral-lifetime unmixed channel images. Scale bar: 10 μm.

### Supplementary information

#### Imaging cardiomyocyte beating

We evaluated compressed FLIM in imaging MacQ-mOrange2 to detect beating in cultured rat heart cardiomyocytes. We transfected cardiomyocytes with plasmid DNA MacQ-mOrange2 and stimulated with acetylcholine. As an example, the induced fluorescence intensity oscillation is shown in **Fig. S6**. We then used compressed FLIM to image the cardiomyocyte beating. **Fig. S7b** shows the fluorescence intensity and lifetime traces of MacQ-mOrange2 sensor expressed in a cultured cardiomyocyte with acetylcholine stimulation imaged at 40 Hz. **Movie 5** records the lifetime dynamics due to cardiomyocyte beating. Representative snapshots at 216 ms, 513 ms, 621 ms, 810 ms, 1080 ms and 1512 ms are shown in **Fig. S7c**. The average relative fluorescence intensity change (Δ*F/F*) and absolute lifetime change (Δτ) in response to one beating event is −1.2% and −0.2 ns. respectively. Finally, to provide a negative control, we imaged MacQ-mOrange2 without acetylcholine stimulation (**Fig. S7d**). Both the fluorescence intensities and lifetimes were stable during the entire time trace, and no beatings were observed.

#### Myocytes transfection

We used Mac mutant plasmid DNA Mac-mOrange2 (#48761, Addgene) to transfect the rat heart cardiomyocytes 10 DIV (H9c2(2-1) ATCC^®^ CRL-1446™), and lipofectamine 2000 (Invitrogen) was used as the transfecting reagent. We imaged the rat heart cardiomyocytes 2 days after transfection. We used acetylcholine to stimulate cardiomyocyte beating. The base medium for this cell line was Dulbecco’s Modified Eagle’s Medium (#30-2002, ATCC). At each stimulation, we removed the media in the plate and pipetted fresh medium containing 100μM acetylcholine to cardiomyocytes^1^.

#### Imaging hematoxylin and eosin (H&E) stained tissue slide

We further validated compressed FLIM in imaging a hematoxylin and eosin (H&E) stained tissue slide. In the sample, we stained connective tissues and extracellular materials using fluorophore eosin. We excited the sample at 515 nm, and imaged the fluorescence using a ×60 1.42 NA oil immersion objective lens (PLAPON 60XO, Olympus). The fluorescence intensity and reconstructed lifetime images are shown in **Fig. S8a and S8b**, respectively. The observed fluorescence lifetime heterogeneity in **Fig. S8b** is attributed to eosin quenching^2^. To validate our results, we imaged the sample using a line-scanning streak camera imaging approach (Methods). The ground-truth image is shown in **Fig. S8c**, matching well with compressed FLIM measurement.

#### Compressed spectral FLIM (sFLIM) imaging

In fluorescence microscopy, multi-target imaging is commonly accomplished by means of spectrally resolved detection and multicolor analysis^3–5^ or time-resolved detection by FLIM^6–7^. However, given a single measurement dimension (spectrum or time), the fluorescence characteristics of co-located fluorophores must differ substantially in order for them to be separated, a fact which limits the number of fluorophores that can be simultaneously imaged. Taking the full advantages of spectral and temporal characteristics of a fluorophore can relieve this requirement and significantly improve the accuracy of unmixing^8^. Herein we expanded the functionality of compressed FLIM to color reproduction and demonstrated compressed spectral FLIM (sFLIM) in imaging multi-target fluorescence samples.

The compressed sFLIM system is shown in **Fig. S9a**. We inserted a bandpass filter in each of two temporally-sheared imaging channels and selectively passed light at given wavelengths: orange (ET590/50m, Chroma) and near infrared (ET690/50m, Chroma). Accordingly, we changed the emission filter of the microscope to a multi-bandpass filter (59007m, Chroma). The resultant temporally-sheared channels record the fluorescence scene filtered at two different wavelengths. To reconstruct the sFLIM image, we separately computed the lifetime images at these two wavelengths using the single-channel TwIST algorithm and registered their images thereafter.

Compared with using lifetime or spectral information alone, the combination of multidimensional fluorescence data can significantly improve the accuracy of fluorophore decomposition. We adopted a previously described algorithm^8,9^ to process the multidimensional data acquired by compressed sFLIM. A probability function *p*(*t, λ*)*dtdλ*, referred to as fluorescence pattern, is constructed to describe how many photons per time interval *dt* and per spectral bandwidth *dλ* are expected from a given fluorophore. At each spatial pixel, the light intensity that compressed sFLIM measures is the superposition of individual fluorescence patterns:

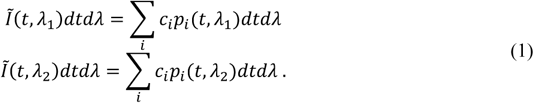

Here, *λ*_1_ and *λ*_2_ are the corresponding peak emission wavelengths in two spectral channels, and *c*_*i*_ denotes the concentration of fluorophore *i*. Decomposition of fluorophores at a given spatial pixel (i.e., calculation of *c*_*i*_) is the solution of the inverse problem of Eq. 1.

The numerical solution of this problem can be found by minimizing the Kullback-Leibler discrepancy *Δ*_*KL*_:

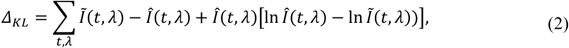

where *Î*(*t, λ*) is the estimation that is calculated by *Î*(*t, λ*) = ∑_*i*_ *ĉ*_*i*_ *p*_*i*_(*t, λ*). A positively constrained solution (i.e., *ĉ*_*i*_> 0 for all *i*) of Eq. 2 can be found via a typical non-negative matrix factorization algorithm^10^. In brief, the algorithm starts with a reasonably estimated positive concentration vector ***ĉ*** and determines the local gradient. During iteration, the concentration vector is updated such a way that it can be expressed as the multiplication with a positively constrained scaling vector ***γ*: *ĉ***_*n*+1_ = ***γĉ***_*n*_.

To demonstrate sFLIM, we imaged a mixture of fluorescent beads with varied lifetimes or emission spectra (6 um diameter orange, F8825; 2 um diameter orange, C16509; 10 um diameter crimson F8831; Thermo Fisher). The fluorescence lifetime and spectral data of these beads are shown in **Fig. S9b** and **Fig. S9c**, respectively. Within a snapshot, we captured the spectrally-resolved lifetime images and show them in **Fig. S9d**. Moreover, we applied the spectral-lifetime unmixing algorithm to the dataset and separated these beads in individual channels as shown in **Fig. S9e**.

